# Expression of Osteopontin in M2 and M4 intrinsically photosensitive retinal ganglion cells

**DOI:** 10.1101/2024.10.31.621275

**Authors:** Leonie Kinder, Moritz Lindner

## Abstract

**Purpose:** Melanopsin-expressing intrinsically photosensitive (ip) retinal ganglion cells (RGC) can be divided into six different subtypes (M1 – M6). Yet, only for some of these subtypes specific markers exist that could be employed to study of the function of individual subtypes. Osteopontin (*Spp1*) marks alpha(α)-RGC, suggesting that, across ipRGC, it would only mark the M4-ipRGC subtype (synonymous to: ON-sustained-αRGC). Recent evidence suggests that osteopontin expression could spread to other ipRGC subtypes. Therefore, this study aims to characterise the expression pattern of osteopontin across ipRGC subtypes.

**Methods:** Single-cell RNA sequencing data from RGC were analysed to identify expression patterns of *Spp1* across ipRGC. Immunohistochemistry (IHC) was performed on retinal cryosections and flatmounts from C57BL/6J mice to characterize the localization of osteopontin across ipRGC. Neurite tracing was employed to study dendritic morphology and identify individual ipRGC subtypes.

**Results:** scRNAseq analysis revealed *Spp1* expression in two distinct clusters of ipRGC. IHC confirmed osteopontin co-localization with Smi-32, an established marker for αRGC, including M4-ipRGCs. Spp1 immunoreactivity was moreover identified in one additional group of ipRGC. By dendritic morphology and stratification those cells were clearly identified as M2-ipRGC.

**Conclusions:** Our findings demonstrate that osteopontin is expressed in both M2- and M4-ipRGC, challenging the notion of osteopontin as a marker exclusively for αRGC. IHC double-labelling for osteopontin and melanopsin provides a novel method to identify and differentiate M2 ipRGC from other subtypes. This will support the study ipRGC physiology in a subtype specific manner and may for instance foster research in the field of optic nerve injury.

## Introduction

In the mammalian retina visual information is processed in more than 40 parallel pathways represented by dedicated subtypes of retinal ganglion cells (RGCs) ^1, 2^. Based on functional commonalities, these subtypes can be classified into different overlappinggroups, like the so-called intrinsically photosensitive retinal ganglion cells (ipRGCs) or the alpha(α)-RGC. Some RGC subtypes exhibit characteristics that align with multiple groups, leading to their classification within more than one of these groups. This is the case, for instance, for the M4-ipRGC subtype, which has been found to be identical with the ON-sustained subtype of αRGCs^3^.

IpRGCs have received major attention after they had been discovered to express the opsin melanopsin (encoded by *Opn4*), which renders them intrinsically light responsive ^4-6^. IpRGCs have since been found to mediate a range of non-image forming responses to light including circadian entrainment or pupil constriction. More recently, it has become clear that these cells also contribute to image forming tasks, for instance by adjusting contrast sensitivity ^7, 8^.This diversity of tasks is not mediated by all ipRGCs equally, rather the different subtypes, termed M1 to M6, appear to be functionally specialized. M1 to M6 ipRGCs can be distinguished based on their *Opn4* expression level, cell morphology and central projection patterns ^8, 9^. Key features are schematically summarized in **Figure 1**.

**Figure 1:**
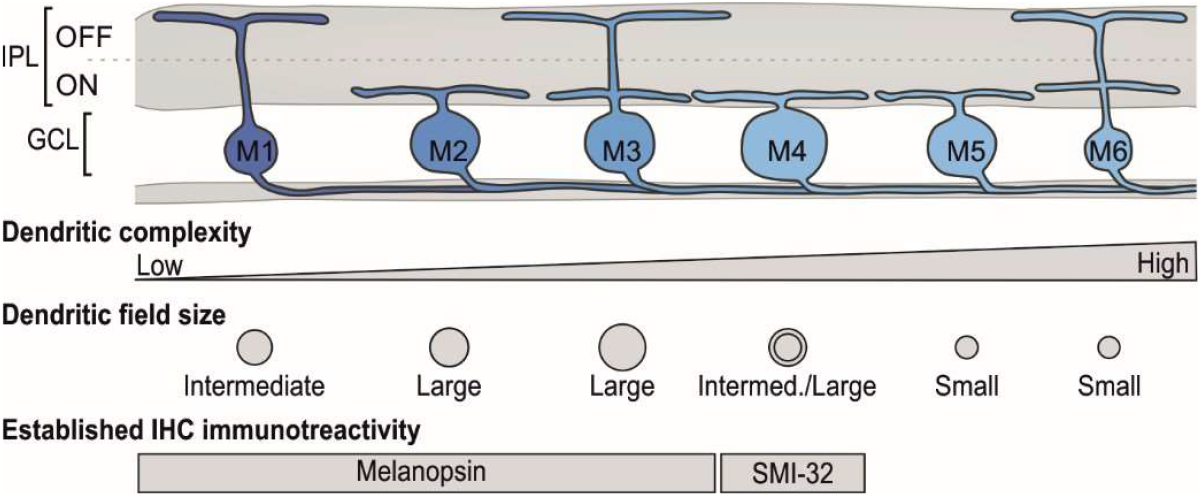
Schematic of the subtypes of Intrinsically Photosensitive Retinal Ganglion Cells (ipRGC) and their structural features. The expression of melanopsin decreases from M1 to M6 (blue). M1 ipRGC are the only subtype stratifying in the OFF lamina of the inner plexiform layer (IPL) while M2, M4 and M5 stratify in the ON lamina and M3 and M6 are bistratifying. Complexity of the dendritic tree increases from the M1 to the M6 subtype. M2/M3 cells have the largest dendritic fields while those of M5 and M6 are small and M1/M4 intermediate. Only M1 – M3 ipRGCs express sufficient melanopsin to be detectable by standard immunohistochemical techniques (For review, see ^7^). GCL: Ganglion Cell Layer, IHC: immunohistochemistry.

Numerous attempts have been made to associate specific ipRGC-mediated physiological functions to individual subtypes. These attempts have been particularly successful for cases where specific immunohistochemical or genetic markers existed to efficiently segregate the subtype of interest from the others. For instance, it could be shown that a specific subpopulation of M1 ipRGC (those lacking *Brn3b* expression) is essential for circadian entrainment ^4, 10^, M4 cells are involved in setting contrast sensitivity ^11^ and M6 likely in pattern vision ^12^. Yet, *Brn3b*^-^ M1, M4, and M6 cells are, to the best of our knowledge, the only ipRGC subtypes for which such markers have been identified, and the functional study of the remaining types still relies on laborious approaches like dye-filling during patch clamp experiments ^13^. Such methods might not enable functional assessment up to a behavioural level.

Thus, identification and careful characterization of potential markers is warranted to enable and accelerate functional characterization of the individual ipRGC subtypes. In this regard, osteopontin (also known as secreted phosphoprotein-1 – Spp1), a multifunctional protein involved in intracellular and extracellular signalling, has caught our attention ^14^. Osteopontin has long been considered a pan-αRGC marker ^15-17^, which would imply that, among the ipRGCs, the M4 subtype should be the only one expressing relevant amounts of osteopontin. In a more recent study, it has been observed that osteopontin immunoreactivity spreads beyond αRGCs, particularly into cells that are also immunopositive for melanopsin (i.e. M1-M3 ipRGCs, **Figure 1**, ^18^). If osteopontin expression was defined to one specific subtype among the M1-M3 ipRGCs, this might open room for specific immunohistochemical or transgenic labelling, that could facilitate segregating the specific functional roles of M1 to M3 cells. This is especially true as mouse lines exist that express cre recombinase under control of the *Spp1* promoter (MGI: 6450712) which would enable flexibly manipulating this subtype. Moreover, understanding how osteopontin expression exactly spreads beyond αRGCs could help to further clarify its potential role in supporting RGC resilience against axonal injury ^15^ and glaucomatous damage ^18, 19^, which a large recent study suggests could be a feature more strongly associated with ipRGCs than to αRGC ^2^.

In this study we therefor examine the expression pattern of osteopontin across ipRGC subtype using a set of transcriptomic and IHC techniques. We find that osteopontin expression outside αRGCs is confined to the M2 subtype of ipRGCs and the combination of osteopontin and melanopsin provides an exclusive IHC marker set for this subtype.

## Results

To explore the expression pattern of osteopontin in ipRGCs, we harnessed a publicly available single-cell RNA sequencing dataset from retinal ganglion cells ^2^. We first identified melanopsin/*Opn4* expressing cells in this dataset (n = 2211) and selected those for further analysis. Unsupervised clustering revealed six transcriptomically distinct populations of ipRGCs (**Figure 2 A**). We then used *Opn4* expression levels as well as expression levels for *Smi32* (coding for Neurofilament heavy chain protein, characteristic for M4 cells ^3^), *Gna14* (coding for the G-protein G_14_α, shown to be functional in M1>M3, ^20^, and *Hcn1/2* (coding for two cyclic nucleotide gated ion-channels, relevant for signal transduction in M4, less so M2 and M3 but not in M1 ^20^, to assign these clusters to the previously described ipRGC subtypes (**Figure 2 B, C**). Probing for *Spp1* expression in those clusters, we found that *Spp1* expression was highest in the cluster presumably representing M2 cells while expression levels detected in the cluster representing M4 cells was markedly lower (log2-fold change = 2.15, p < 0.001, **Figure 2 C, D, E**)

**Figure 2.**
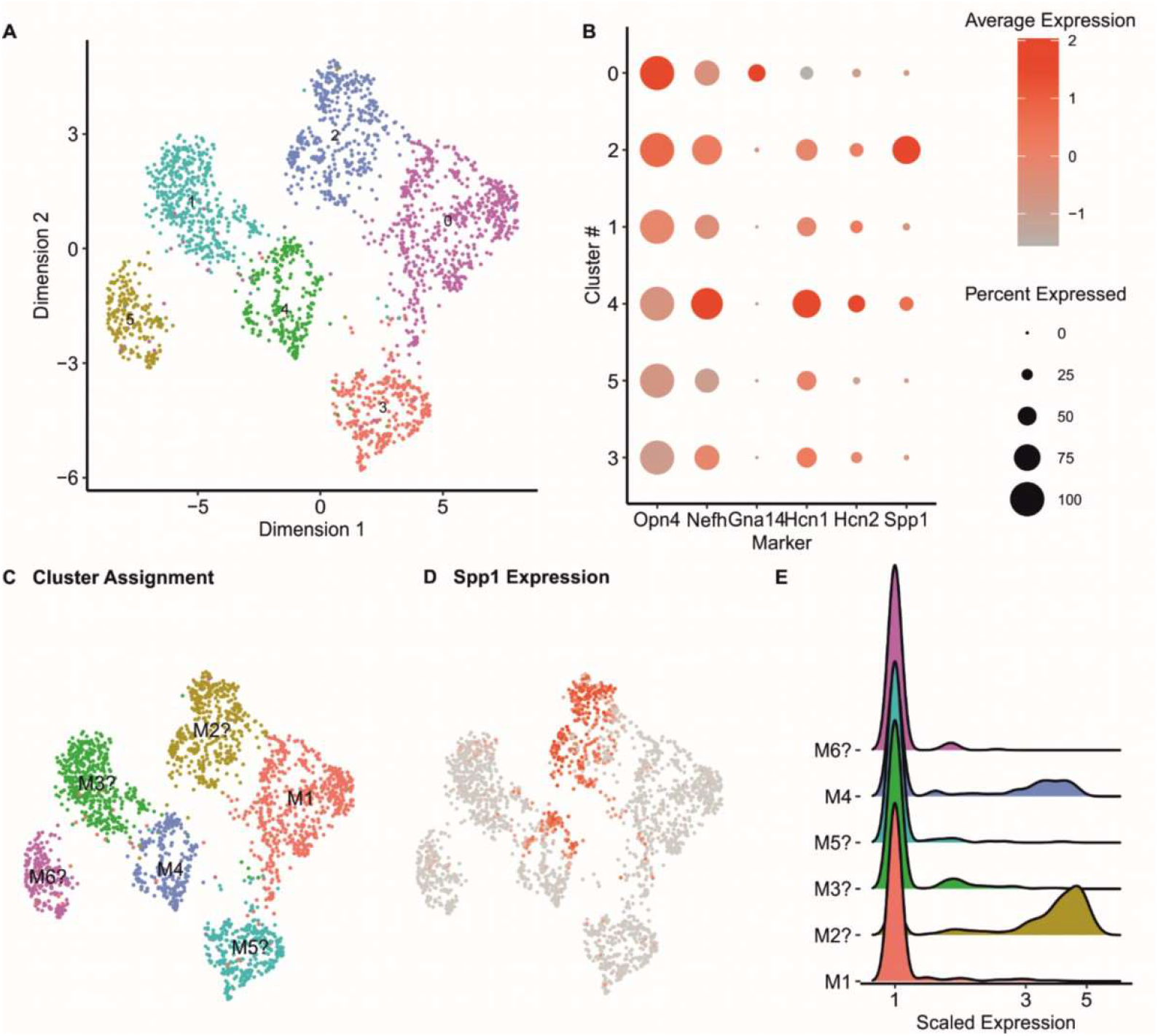
Transcriptomic expression patterns of Spp1 across ipRGC subtypes. Re-analysis of the Opn4-expressing cells from a publicly available single-cell RNA-sequencing dataset ^2^. **(A)** Clustering and UMAP visualization after dimensionality reduction reveals six independent clusters of Opn4-expressing cells (#0-5). **(B)** Expression patterns of established markers were used to grossly annotate ipRGC subtypes ^3, 16, 21-23^. **(C, D)** UMAP visualization as in (A), but with annotated clusters (C) and colour-coded expression of Spp1 (D, scale bar as in B). **(E)** Ridgeline plot showing the distribution of Spp1 expression in the different clusters.

To probe whether these transcript level observations would be reflected on protein level, we continued our analysis performing IHC staining on retinal cryosections. We performed triple labelling for osteopontin together with Smi-32 as bona-fide pan-marker for αRGCs, including M4 ^3, 16, 21, 22^ and melanopsin to identify M1-M3-ipRGCs. The results showed the previously reported co-localization of Smi-32 and osteopontin in the ganglion cell layer. These Smi-32^+^/osteopontin^+^ cells appeared to have rather large somas, consistent with the assumption that these would be αRGCs (**Figure 3 A, B**). We, however, observed also a smaller fraction of cells that showed osteopontin immunoreactivity while being immunonegative for Smi-32. The vast majority of these cells were immunopositive for melanopsin (**Figure 3 C, D**), suggesting that these must be M1, M2 or M3 ipRGCs. While most of them had their soma in the ganglion cell layer, we found one cell with a rather weak osteopontin labelling residing in the inner nuclear layer. Of note, a small portion of osteopontin-positive cells was negative for both, Smi32 and melanopsin, indicating the presence of other RGC types expressing also osteopontin.

**Figure 3.**
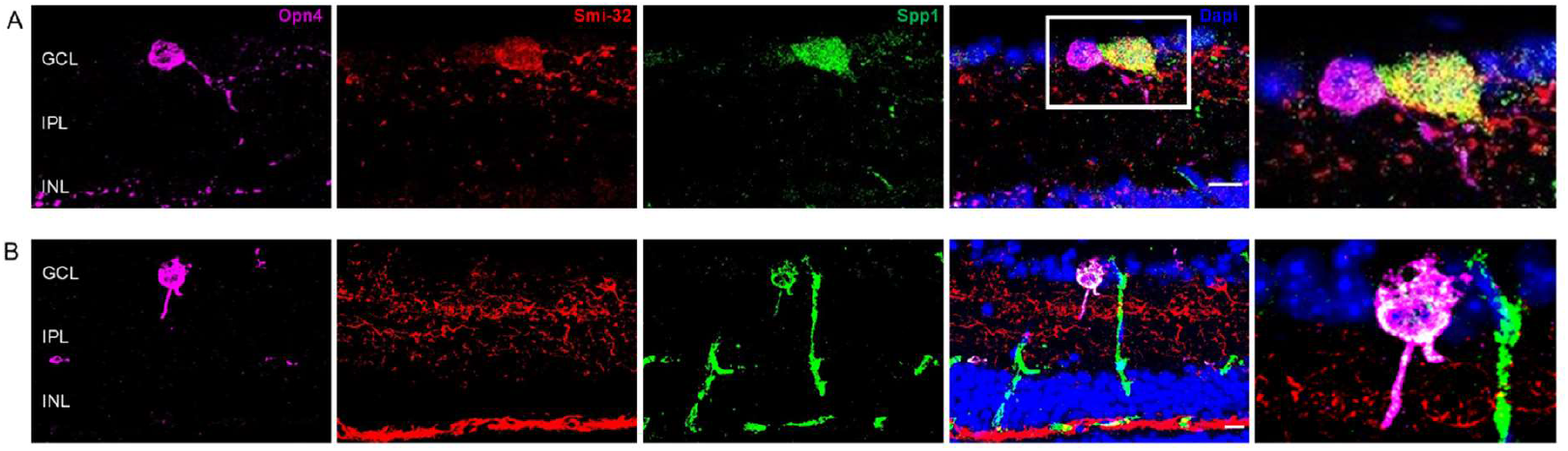
Osteopontin immunoreactivity in αRGC and a subset of melanopsin-immunopositive ipRGCs. Confocal micrographs of retinal cryosections immunostained for melanopsin (=Opn4, magenta), neurofilament-heavy chain protein (=Smi-32, red) and osteopontin (=Spp1, green). ***(A)*** Representative example of a cell where Spp1 immunoreactivity co-localizes with Smi-32, but not with Opn4. (***B)*** Representative example of a cell where Spp1 immunoreactivity co-localizes with Opn4Smi-32, but not with Smi-32, most likely representing an M1, M2 or M3-ipRGC. Blood vessels showing up in the Spp1-channel (green) due to murine origin of the osteopontin antibody. GCL =ganglion cell layer, IPL=inner plexiform layer, INL =inner nuclear layer. *Blue channel: DAPI. Scale bar 10 μm*.

We then turned to retinal flatmount staining. This allowed us to assess much larger areas of the retina and thereby quantify the relative abundance of osteopontin expressing M1-M3 ipRGCs. Performing colabelling for melanopsin and osteopontin we found 147 ± 29 melanopsin positive (M1-M3) ipRGCs and a total number of 143 ± 22 osteopontin positive cells. 78 ± 23 co-labelled for melanopsin and osteopontin, thus representing roughly 50 % of all M1-M3 ipRGCs. Performing confocal volume scans on these retinal flatmounts moreover allowed us to trace the dendritic arbor of individual cells, enabling us to determine dendritic stratification, which characteristically distinguishes between M1, M2 and M3 subtypes. Per flatmount stained, we traced the dendritic tree of one index cell (osteopontin^+^/melanopsin^+^) and one neighbouring control cell (osteopontin^-^/melanopsin^+^, n = 6 each). All index cells were monostratifying in the ON-sublamina of the IPL exclusively, suggesting they were M2 ipRGCs. By contrast control cells were either monostratifying in the OFF-layer or bi-stratifying in ON-and OFF-layer (**Figure 4 and 5A**), therefore most likely representing the ipRGC subtypes M1 (OFF) and M3 (ON-OFF). In agreement with these findings, we observed that the index cells had a large dendritic field diameter of 253.91 ± 20.3 μm and many (20 ± 3) dendritic branching points, while the control cells had a smaller dendritic field diameter of 243.47 ± 44.6 μm and less (12 ± 3) branching points (**Figure 5 B and C**). Also, in accordance with these findings the mean intensity of the melanopsin immunofluorescence measured over the somata of index cells was significantly lower as compared to that of control cells. As for the osteopontin-intensity, there was moderately but significantly weaker signal in the osteopontin^+^/melanopsin^+^ cells compared to cells that only expressed osteopontin but no melanopsin (i.e. mostly αRGC; relative fluorescence intensity in osteopontin^+^/melanopsin^+^ was 73 [inter quartile range: 65 - 79] % of that observed in osteopontin^+^/melanopsin^-^ cells; p = 0.032).

**Figure 4.**
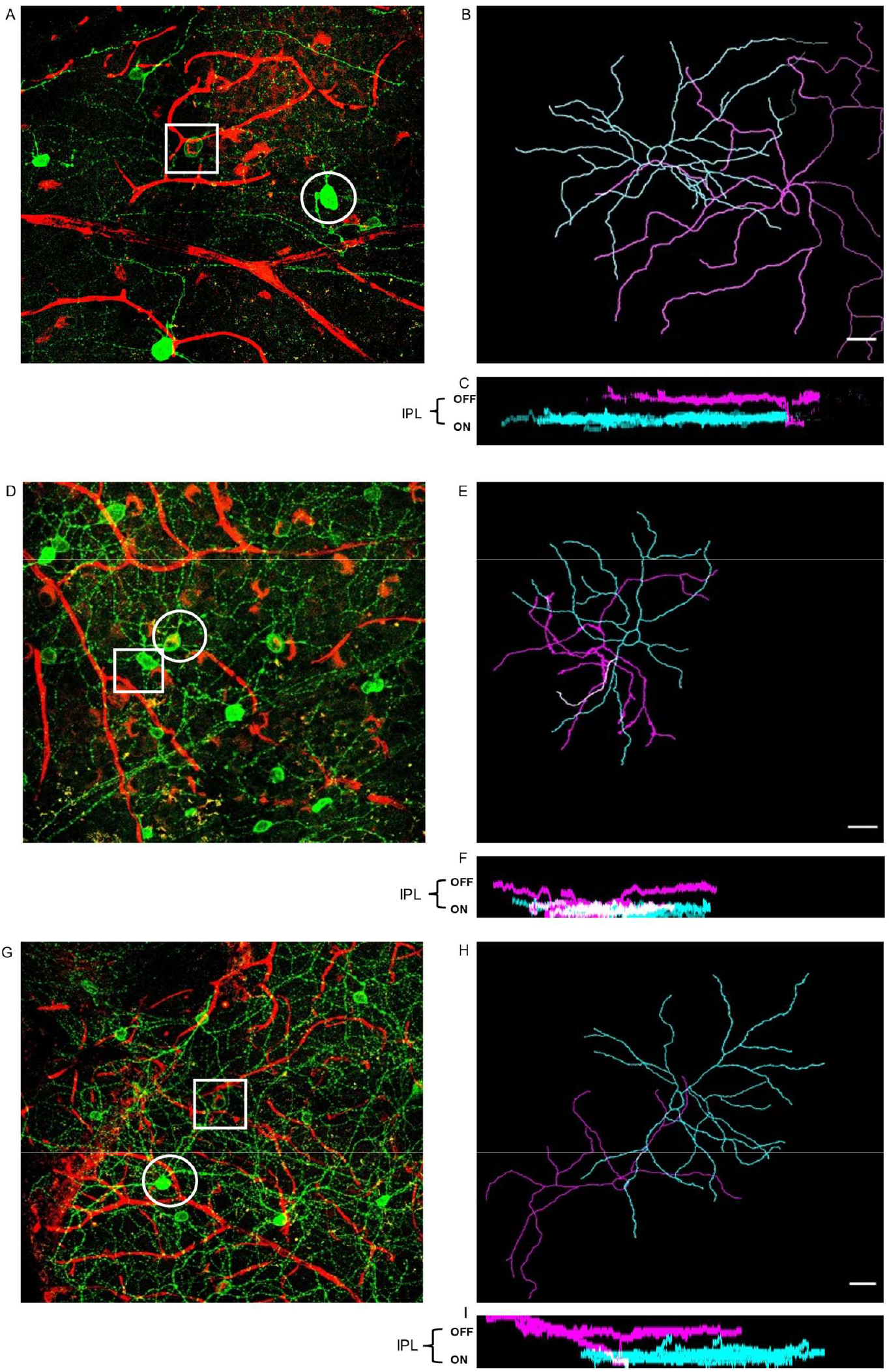
Osteopontin immunoreactive ipRGCs structurally resemble M2 cells. (*A, D, G,)* Exemplary Z-projections of confocal volume-scans from murine retinal flatmounts immunolabeled for osteopontin (=Spp1, red) and melanopsin (=Opn4, green). Spp1-positive cell (,,index-cell”, square) next to Spp1-negative ipRGC (,,control-cell”, circle) (***B, E, H)*** Outlines of neurite-traced neurons: Spp1-positive index cell (cyan) next to Spp1-negative control cell (magenta).(***C, F, I)*** Orthogonal representation of the traced cells showing the index cells monostratifying in the ON-sublayer of the IPL and control cells either monostratifiying in the OFF-layer **(C, I)** or bistratifying in ON- and OFF-sublayer of IPL **(F)**. Blood vessels showing up in the Spp1-channel (green) due to murine origin of the osteopontin antibody. Scale bar 30μm.

**Fig. 5.**
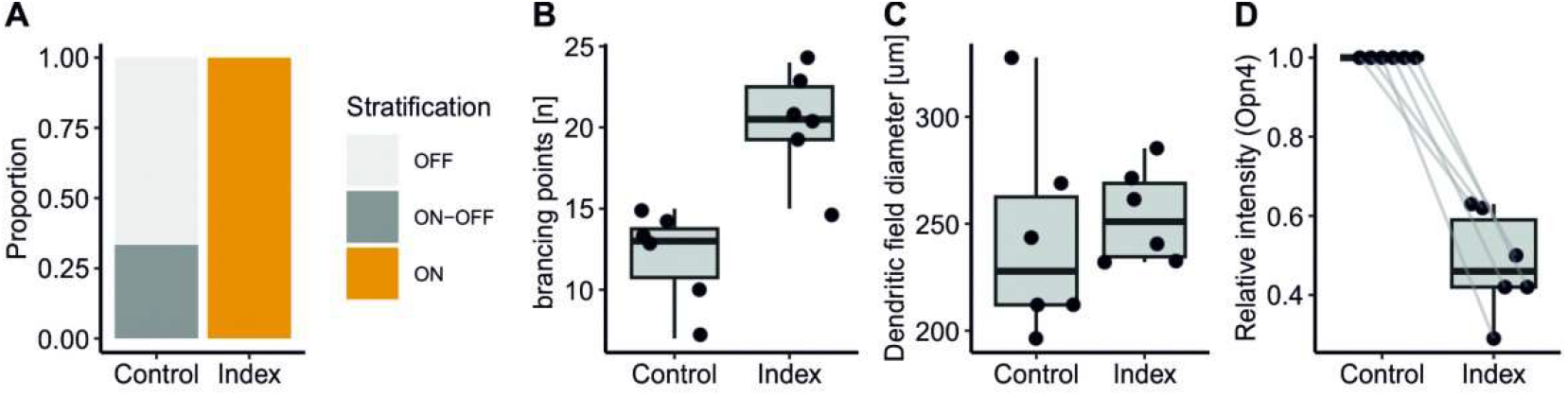
Statistical analysis of morphological characteristics found in traced index and control cells. (*A)* Stratification patterns of index and control-cells, showing index cells stratifying exclusively in the ON-sublamina of the IPL, while traced control cells were monostratifying in the OFF-sublayer (66,66%) and bistratifying in ON- and OFF-layer (33,33%). (***B)*** Average number of branching points of index and control-cells showing a more complex dendritic arborization of our index cells along with a larger dendritic field diameter as shown in (***C)***. (***D)*** Relative pixel intensity of Opn4 of index cell in relation to respective control cells. Index cells show a significantly lower Opn4-intensity.

## Discussion

In the present work, we provide an in-depth characterization of the expression pattern of osteopontin across ipRGC subtypes. Using transcriptomic and immunohistochemical techniques, we demonstrate that not only M4 – as previously thought – but also M2 ipRGCs express osteopontin. Together with melanopsin, osteopontin assembles a marker pair that allows for efficient identification of M2 ipRGCs in immunohistochemical experiments. This will facilitate future studies aiming to dissect the functional roles of individual ipRGC subtypes.

To draw this conclusion, we commenced by analysing the ipRGCs contained in a publicly available RNAseq dataset ^2^ and we found Osteopontin expression on two distinct clusters. Judged by the expression of αRGC marker Smi-32 and signal transduction genes differentially operative in M1 to M6 ipRGCs ^11, 20, 23^, these clusters resembled M2 and M4 cells, respectively. Indeed, using immunohistochemical approaches we could first confirm the existence of a Smi-32-negative, i.e. none-M4 population of cells that was mostly immunopositive for melanopsin. We could then show that this osteopontin^+^/melanopsin^+^ population exhibited structural features which are characteristic for the M2 subtype, most prominently, the ON-lamina stratification of their dendritic tree ^9^. Consistently, osteopontin^+^/melanopsin^+^ cells had a larger, more complex, dendritic tree and expressed lower levels of melanopsin as compared to osteopontin negative ipRGCs. Moreover, making up approx. 50% of all Opn4^+^ cells, this proportion nicely matches that expected for M2 – but was far too high for the alternative but rare M3 subtype ^24, 25^.

As mentioned above, osteopontin expression has been previously considered to be αRGC-exclusive and previous studies have thus used osteopontin as a marker (or genetic selector) for this cell type ^15, 17, 26^. The present study, as well as the results by Zhao et al. ^18^ now show that this is indeed not fully precise. As a whole, αRGC clearly outnumber M2-ipRGCs. However, if selecting αRGC by osteopontin expression, the systematic inclusion of M2 cells would induce an unwanted bias, that could be more or less relevant depending on the specific experimental question.

In our immunohistochemical experiments we also noticed a small portion of osteopontin-positive cells, that neither expresses Smi-32 nor melanopsin. In theory, these cells could be a yet entirely different type of RGC or any of the two other ipRGC subtypes (besides M4) with melanopsin expression levels too low for immunohistochemical detection, i.e. either M5 or M6. From our transcriptomic analyses, it appears likely that these cells were rather not M5 or M6, as the clusters representing these subtypes yielded expression level of *Spp1* that were virtually zero.

One additional observation we made is that on transcriptomic level *Spp1* expression was even higher in the M2 cluster than in the M4 cluster. We were not able to test if that observation was preserved to protein level, as we did not distinguish between M4 and the other Smi-32-positive αRGCs in IHC. Yet, based on the transcriptomic results alone, it would seem that in fact M2 cells might be those with the strongest osteopontin expression among the ipRGCs. Notably, this does not mean that M2 osteopontin levels would exceed those of all other RGC types in general, rather our immunostaining experiments indicate that osteopontin levels are lower in M2 cells than in the overall αRGC population (i.e. osteopontin expression M4<M2<non-M4-αRGC).

In this study, we aimed to characterize the expression patterns of osteopontin across ipRGCs, considering osteopontin as a cellular marker, while its functional role in these cells has not been our focus. Interestingly, osteopontin had been suggested to render RGCs resilient to (axonal) injury or other damage. Possibly this concept might have been led by the strong overlap between the cells belonging to the group of αRGC and those expressing osteopontin. Indeed, it has been shown that osteopontin supports RGC survival, possibly in an interplay with Insulin-like growth factor receptor 1. Yet, while supportive, it is clearly not essential for it ^15^. On the other end, more recent results based on data-driven techniques demonstrate that resilience is predominantly a feature of ipRGCs rather than αRGC and while transcriptomic determinants for RGC resilience can be identified, *Spp1* is not one of those genes with the strongest effect ^2^. Still, the refined characterization of osteopontin expression across ipRGC subtypes provided herein, will support attempts to identify therapeutically usable factors supporting RGC survival and resilience, like promoted by the RReSTORe consortium ^27^.

The approach chosen in this study, starting off from subsetting ipRGCs by *Opn4* expression from a large transcriptomic dataset ^2^ has enabled us to robustly cluster this relatively rare set or retinal neurons. While we have used this approach to assess the expression patterns of *Spp1*/osteopontin herein, it can also serve as a more general approach for identification of ipRCG subtype markers in future studies. For such purposes other large single cell datasets like that published by Rheaume et al ^28^ may serve as a reference to narrow down the list of candidate markers. For characterizing the expression pattern of one particular marker, in this study we performed in-depth structural characterization of osteopontin^+^/melanopsin^+^ (and osteopontin^-^/melanopsin^+^ control cells) for a relatively small number of samples using manual neurite tracing. While the small-ish sample size might be somewhat of a limitation to this study, observing the characteristic M2 ON-sublamina stratification in 100% of the index and 0% of the control cells, together with the observation that all other morphometric measures fitted well into this hypothesis, does provide a high degree of confidence that our findings were indeed robust.

In conclusion, in the present study we could demonstrate that osteopontin is expressed in two distinct subtypes of ipRGCs, namely M2 and M4 cells. The combination of osteopontin and melanopsin immunoreactivity thereby provides an exclusive marker set for M2 ipRGCs.

## Material and methods

### Animals

C57BL/6J mice were purchased from Charles River Laboratories (Sulzfeld, Germany). Handling of the mice and tissue collection was performed in accordance with the federal and institutional guidelines.

### Tissue preparation

For histological analysis, mice were culled by decapitation under isoflurane anaesthesia and tissue was collected and processed following to in-house standard procedures ^29^. Eyes were punctured with a fine gauge needle, fixed in 4% methanol-free paraformaldehyde (Thermo Fisher, Waltham, MA, USA) in phosphate buffered saline (PBS) for 30 min (cryosections) or 24 h (flatmounts), and then transferred to 30% sucrose in H_2_O. For preparation of retinal cryosections eyes were embedded into Optimal Cutting Temperature (OCT) medium (VWR, Lutterworth, UK) and stored at -80 ºC until further processing. 18 μm tissue sections were prepared using a CM1850 Cryotome (Leica, Wetzlar, Germany). For retinal flat mount preparation, the anterior segment of the fixed eye was removed cutting posterior to the ora serrata, and the retina was mobilized by gently lifting it off from the underlying RPE with sclera. Four releasing incisions were made into the retina before storing it in PBS until further use.

### Immunohistochemistry

Fluorescent immunolabelling was performed using standard techniques as previously described ^29^. Briefly, for retinal cryosections were permeabilized in phosphate buffered saline (PBS) with 0.2% Triton-X (PBSTX-0.2). Blocking was performed in PBSTX-0.2 with 10% normal donkey or goat serum (Sigma Aldrich, St. Louis, USA). Sections were then incubated with primary antibodies for 24 h at 4 °C and after washing with 0,1% Tween in PBS incubated with secondary antibodies for another 2 hours at room temperature. Sections were mounted with Prolong Gold antifade media (Life Technologies / Thermo Fisher Scientific, Carlsbad, USA). All antibodies were diluted in PBSTX-0.2 containing 2.5% normal donkey serum.

For flat mount staining, 1% Triton-X in PBS (PBSTX-1) was used for permeabilization. Blocking was performed in PBSTX-1 with 10% normal donkey or goat serum (Sigma Aldrich). Primary antibody incubation was performed over three days at room temperature, followed by incubation with the second antibody overnight at room temperature. Washing steps were performed in 0,1% Tween in PBS. Antibody soles were made up in PBSTX-1 with 2,5% normal donkey/goat serum. A list of all primary and secondary antibodies used in this study is given in **Table 1**.

**Table 1:**
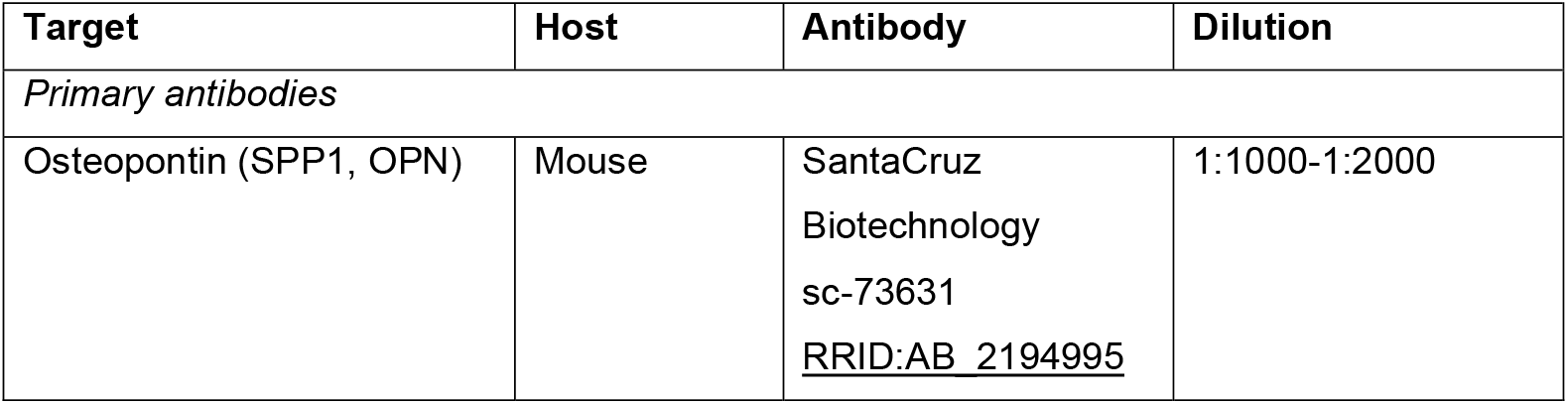
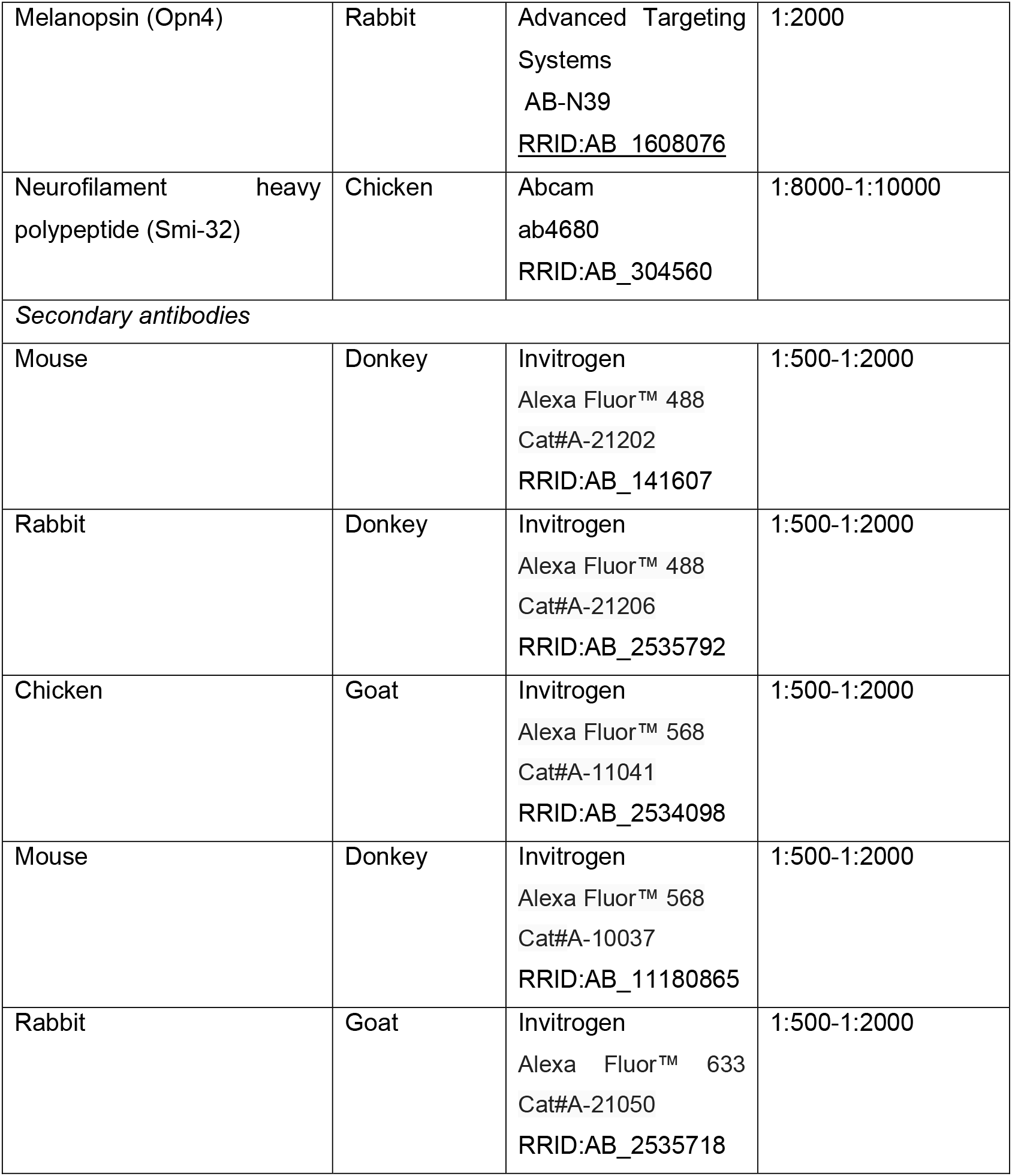
List of primary and secondary antibodies used in this study.

| Target | Host | Antibody | Dilution |
| --- | --- | --- | --- |
| *Primary antibodies* |  |  |  |
| Osteopontin (SPP1, OPN) | Mouse | SantaCruz<br>Biotechnology<br>sc-73631<br>RRID:AB_2194995 | 1:1000-1:2000 |
| Melanopsin (Opn4) | Rabbit | Advanced Targeting Systems<br>AB-N39<br><u>RRID:AB_1608076</u> | 1:2000 |
| Neurofilament heavy polypeptide (Smi-32) | Chicken | Abcam<br>ab4680<br>RRID:AB_304560 | 1:8000-1:10000 |
| *Secondary antibodies* |  |  |  |
| Mouse | Donkey | Invitrogen<br>Alexa Fluor™ 488<br>Cat#A-21202<br>RRID:AB_141607 | 1:500-1:2000 |
| Rabbit | Donkey | Invitrogen<br>Alexa Fluor™ 488<br>Cat#A-21206<br>RRID:AB_2535792 | 1:500-1:2000 |
| Chicken | Goat | Invitrogen<br>Alexa Fluor™ 568<br>Cat#A-11041<br>RRID:AB_2534098 | 1:500-1:2000 |
| Mouse | Donkey | Invitrogen<br>Alexa Fluor™ 568<br>Cat#A-10037<br>RRID:AB_11180865 | 1:500-1:2000 |
| Rabbit | Goat | Invitrogen<br>Alexa Fluor™ 633<br>Cat#A-21050<br>RRID:AB_2535718 | 1:500-1:2000 |

### Image acquisition and analysis

Fluorescence images were acquired using an upright LSM 710 laser scanning confocal microscope (Zeiss, Oberkochen, Germany) equipped with an 40x oil objective (NA = 1.3). Laser lines for excitation were 405 nm, 488 nm, 561 nm and 633 nm with emissions collected between 440–480, 505–550, 580–625 nm and >640 nm, respectively. Individual channels were collected sequentially. Image post-processing was restricted to global enhancement of brightness and contrast, as well as cropping, downscaling and sub-selecting of fluorescent channels as required, Post processing was performed using ImageJ/FIJI ^30^. For retinal flatmounts, Z-stacks (slice interval 0,45μm, average depth: 16,87±1,36 μm) were acquired from four adjacent areas and were stitched together using FIJIs 3D-stitching tool ^31^ to ensure coverage of the entire dendritic field of any neuron in the centre of the stitched image. The ImageJ plugin Simple Neurite Tracer was used for semi-automatic neuron tracing ^32^. Per stitched composite image, at least one index (osteopontin^+^/melanopsin^+^) and one control cell (osteopontin^-^/melanopsin^+^) was traced and signal intensity, number of branching points and dendritic field area were quantified. Signal intensity was quantified as the mean pixel intensity in a 3D region of interest covering the soma of the cell under investigation. As neither laser stimulus intensity nor acquisition parameters were normalized, intensity values were calculated as relative to a control cell on the same image.

### Bioinformatics

To characterize the transcriptomic profile of the individual ipRGC subtypes, a publicly available single cell RNA sequencing dataset ^2^ was analysed. Gene count matrices obtained from the Gene Expression Omnibus (GEO) server, www.ncbi.nlm.nih.gov/geo/ (accession number GSE137863). Processing and analysing of the dataset were performed using the R software environment (version 3.5.1) and the Seurat package (version 5). ^33, 34^ Clustering was performed after dimensionality reduction by Seurat’s inbuilt shared nearest neighbour modularity optimization-based clustering algorithm.

## Acknowledgements

We thank Prof Burkhard Schütz and the Dept. of Anatomy for providing access to their cryotome.

## Declarations

### Author Contributions

*Participated in research design: LK, ML*

*Conducted experiments: LK*

*Performed data analysis: LK, ML*

*Wrote or contributed to the writing of the manuscript:* LK, *ML*

### Data Availability

The datasets generated during and/or analysed during the current study are available from the corresponding author on reasonable request.

### Ethics approval

This study did not involve human participants. Animal work was performed with approval of the relevant authorities and in accordance with the institutional Ethics Guidelines of Animal Care. Further details are provided in the Methods section.

